# Phylogenomic subsampling and the search for phylogenetically reliable loci

**DOI:** 10.1101/2021.02.13.431075

**Authors:** Nicolás Mongiardino Koch

## Abstract

Phylogenomic subsampling is a procedure by which small sets of loci are selected from large genome-scale datasets and used for phylogenetic inference. This step is often motivated by either computational limitations associated with the use of complex inference methods, or as a means of testing the robustness of phylogenetic results by discarding loci that are deemed potentially misleading. Although many alternative methods of phylogenomic subsampling have been proposed, little effort has gone into comparing their behavior across different datasets. Here, I calculate multiple gene properties for a range of phylogenomic datasets spanning animal, fungal and plant clades, uncovering a remarkable predictability in their patterns of covariance. I also show how these patterns provide a means for ordering loci by both their rate of evolution and their relative phylogenetic usefulness. This method of retrieving phylogenetically useful loci is found to be among the top performing when compared to alternative subsampling protocols. Relatively common approaches such as minimizing potential sources of systematic bias or increasing the clock-likeness of the data are found to fare worse than selecting loci at random. Likewise, the general utility of rate-based subsampling is found to be limited: loci evolving at both low and high rates are among the least effective, and even those evolving at optimal rates can still widely differ in usefulness. This study shows that many common subsampling approaches introduce unintended effects in off-target gene properties, and proposes an alternative multivariate method that simultaneously optimizes phylogenetic signal while controlling for known sources of bias.

## Introduction

During the last decades, molecular datasets composed of thousands of genes have become common. Although a few phylogenetic questions have remained uncertain even in the face of such large datasets (King and Rokas 2017; Smith, et al. 2020), phylogenomics has greatly improved our understanding of the structure of the tree of life (Dunn, et al. 2008; Spang, et al. 2015; Burki, et al. 2020), the timing of origin of major clades (dos Reis, et al. 2012), and the changes in genomic architecture associated with key evolutionary transitions (Paps and Holland 2018; Fernández and Gabaldón 2020). At the same time, the analysis of phylogenomic datasets has posed numerous novel challenges. These range from a high prevalence of genes whose evolutionary histories deviate from that of the group of species under study (such as results from events of paralogy, incomplete lineage sorting and hybridization, among others), to an accumulation of non-phylogenetic signals as a product of heterogeneities in evolutionary processes. While many of these issues can be alleviated by implementing more complex models of molecular evolution, computational limitations often preclude their use with entire phylogenomic datasets (Simion, et al. 2020).

Phylogenomic subsampling is a common procedure to alleviate these issues (Chen, et al. 2015; Edwards 2016; Simmons, et al. 2016; Molloy and Warnow 2018; Mongiardino Koch 2019). By focusing on a small fraction of genes that are considered more reliable, contentious or unstable nodes can be tested, and the effects of potentially confounding factors such as missing data and saturation can be disentangled (Fernández, et al. 2014; Sharma, et al. 2014; Borowiec, et al. 2015; Kocot, et al. 2017; Mongiardino Koch, et al. 2018; Stiller, et al. 2020). Smaller datasets are also amenable to analysis using more complex and computationally demanding approaches, including inference under site heterogenous and multispecies coalescent models (Whelan, et al. 2015; Thawornwattana, et al. 2018; Ballesteros, et al. 2019; Marlétaz, et al. 2019). Phylogenomic subsampling can therefore reduce heterogeneities in the dataset and improve model fit, producing results that are often preferred. The same logic applies to divergence-time estimation, where subsampling can be used to both alleviate computational burden and produce more accurate results (Dornburg, et al. 2014; Smith, et al. 2018; Carruthers, et al. 2020; Mongiardino Koch and Thompson 2020).

Given these benefits, multiple subsampling protocols have been proposed. While sharing a common goal of retrieving phylogenetically reliable loci (throughout, used interchangeably with genes), they have often employed—and sought to optimize—entirely different criteria. These can either be a measure of information quantity, such as the length of the alignment or its proportion of missing data/occupancy (e.g., Hosner, et al. 2016; Foley, et al. 2019), or a variable reflecting information quality. Among the latter, common approaches include the selection of loci with high levels of phylogenetic signal (e.g., Salichos and Rokas 2013) and the removal of those potentially affected by systematic biases (e.g., Nesnidal, et al. 2010). However, multiple sources of bias are known (Kapli, et al. 2021) and different proxies for signal have been employed (Salichos and Rokas 2013; Salichos, et al. 2014; Arcila, et al. 2017; Philippe, et al. 2019; Vankan, et al. 2020), and the downstream consequences of choosing among these are largely unknown. This is further complicated by the fact that sources of bias and proxies for signal can be strongly correlated (Mongiardino Koch and Thompson 2020), such that the optimization of either dimension individually modifies the other in potentially unintended ways. As a consequence, it remains unclear if these alternatives (retaining “good” genes vs. discarding “bad” ones) converge on a similar pool of reliable loci, and if not, whether one systematically outperforms the other.

Ultimately, levels of both signal and noise are manifestations of underlying differences in rates of evolution. Rate-based subsampling is therefore also common, but there seems to be little consensus on how it should be implemented: studies have variously supported the use of molecular data that evolve at fast, intermediate or slow rates, as well as the generation of partitions with homogenous rates (e.g., Cummins and McInerney 2011; Rota-Stabelli, et al. 2011; Fernández, et al. 2014; Sharma, et al. 2014; Telford, et al. 2014; Sharma, et al. 2015; Streicher, et al. 2018; Rangel and Fournier 2019; Evangelista, et al. 2021; Li, et al. 2021). These studies have also relied on different types of rate estimates—including tree- and alignment-based metrics of substitution rates, measures of character similarity and compatibility, and proportions of variable/informative sites—as well as different units of measurement (sites or loci). Furthermore, the discovery of appropriate rates of evolution can be complicated by heterogeneities among sites and lineages that are often not accounted for (Dornburg, et al. 2019). An alternative method involves using some notion of the relationships among the taxa under study (including topology and branch lengths in units of time) to predict the likely behavior of data evolving under differing rates (Townsend 2007; Townsend, et al. 2012; Su and Townsend 2015). This approach, termed phylogenetic informativeness (PI), can be used to quantify the expected probabilities of sites contributing towards correctly or incorrectly resolving a given quartet, guiding the discovery of particularly useful genes (e.g., Alda, et al. 2019; Bellot, et al. 2020).

While many studies have optimized just one of these properties, others have devised complicated subsampling schemes intended to find loci that satisfy a number of requisites. In the majority of cases, this is performed by iteratively removing data based on a number of rules (e.g., Fernández, et al. 2014; Sharma, et al. 2015; Whelan, et al. 2015). To some extent, this approach can be used to test the effect of individual gene properties on phylogenetic reconstruction, as well as progressively narrow in on a small set of loci that satisfy multiple criteria. However, the final results depend on the order in which properties are evaluated and the thresholds enforced, decisions that are difficult to justify (if not entirely arbitrary). A handful of studies (Borowiec, et al. 2015; Kocot, et al. 2017; Mongiardino Koch and Thompson 2020) have therefore selected loci that simultaneously satisfy a number of conditions. In the case of Mongiardino Koch and Thompson (2020), subsampling was not performed directly on the variables measured but on principal component axes derived from these. This approach produced axes capturing differences in rate of evolution and overall phylogenetic usefulness along which loci could be sorted. Whether major axes of variation in other phylogenomic datasets can be interpreted in similar ways remains unknown.

Several recent studies have explored a number of these gene properties in an attempt to discover reliable predictors of the phylogenetic performance of loci (Aguileta, et al. 2008; Doyle, et al. 2015; Shen, et al. 2016; Brown and Thomson 2017; Kuang, et al. 2018; Burbrink, et al. 2020; Vankan, et al. 2020; Evangelista, et al. 2021). Their recommendations have often differed, raising the possibility that a universal predictor might not exist. They have also invariably focused on correlating alternative properties with measures of topological distance or clade support, without actually evaluating the performance of subsampled datasets composed of multiple loci. In this study, I calculate numerous gene properties across 18 phylogenomic datasets, representing diverse clades whose evolutionary histories began anytime between the Middle Cambrian and the Late Cretaceous (Table 1). With these data I explore the existence of universal patterns of covariance between gene properties, and test whether such patterns capture useful information regarding the evolutionary history of loci. I then analyze the success of alternative subsampling strategies in finding phylogenetically reliable datasets of small sizes.

**Table 1:** Phylogenomic datasets employed. Age constitutes the inferred date of the last common ancestor of the ingroup (in million years, Ma) as estimated by the same study. Number of taxa corresponds only to ingroup taxa, number of loci to those for which all properties could be estimated (see Methods); these and other numbers differ from those reported in the origin studies.

| Dataset | Age<br>(Ma) | Number<br>of taxa | Number<br>of loci | Occupancy<br>(%) | Mean locus<br>length |
| --- | --- | --- | --- | --- | --- |
| Actinopterygii (Hughes, et al. 2018) | 376.3 | 302 | 1035 | 81.2 | 167.1 |
| Araneae (Fernández, et al. 2018) | 366.1 | 160 | 1114 | 64.2 | 218.8 |
| Aspergillacea (Steenwyk, et al. 2019) | 117.4 | 81 | 1660 | 97.5 | 633.8 |
| Blattodea (Evangelista, et al. 2019) | 206.7 | 45 | 2556 | 82.1 | 374.4 |
| Echinoidea (Mongiardino Koch and Thompson 2020) | 265.0 | 34 | 2356 | 71.6 | 257.1 |
| Gnathostomata (Irisarri, et al. 2017) | 457.6 | 100 | 4543 | 81.6 | 430.4 |
| Heliozelidae (Milla, et al. 2020) | 84.0 | 38 | 1040 | 92.2 | 271.4 |
| Hemipteroids (Johnson, et al. 2018) | 420.3 | 171 | 2225 | 90.6 | 771.0 |
| Hexapoda (Misof, et al. 2014) | 479.1 | 134 | 1467 | 94.7 | 869.5 |
| Hymenoptera (Peters, et al. 2017) | 281.0 | 169 | 2665 | 84.8 | 647.6 |
| Lepidoptera (Kawahara, et al. 2019) | 299.5 | 186 | 2021 | 88.8 | 359.4 |
| Monilophytes (Shen, et al. 2018a) | 321.1 | 69 | 2357 | 89.5 | 284.3 |
| Myriapoda (Fernández, et al. 2016) | 504.4 | 40 | 1942 | 82.2 | 297.1 |
| Opiliones (Fernández, et al. 2017) | 414.2 | 54 | 1288 | 63.2 | 265.7 |
| Phasmatodea (Simon, et al. 2019) | 121.8 | 38 | 1022 | 88.6 | 772.3 |
| Pseudoscorpiones (Benavides, et al. 2019) | 337.5 | 41 | 2110 | 63.2 | 376.1 |
| Saccharomycotina (Shen, et al. 2018b) | 404.0 | 332 | 2348 | 88.1 | 464.6 |
| Scorpiones (Sharma, et al. 2018) | 381.3 | 30 | 1462 | 86.6 | 226.3 |

## Results

Datasets analyzed were coded as amino acids and were modified only by removing loci with less than 50% occupancy. Time-calibrated species trees were also obtained from the corresponding studies. Gene trees were inferred using ParGenes v. 1.0.1 (Morel, et al. 2018) under optimal models, and 100 replicates of non-parametric bootstrap (BS) were used to calculate node support. Site-wise rates of evolution were estimated using the empirical Bayes method implemented in Rate4Site (Mayrose, et al. 2004). All other analyses were performed in the R statistical environment (R Core Team 2019) using custom scripts. This included the estimation of 15 gene properties: 1) alignment length; 2) proportion of missing data; 3) level of occupancy; 4) proportion of variable sites; 5) total tree length (i.e. sum of all branches); 6) level of treeness (i.e., the fraction of tree length on internal branches; Phillips and Penny 2003); 7) average pair-wise patristic distance between terminals, a proxy for sensitivity to long-branch attraction (Struck 2014); 8) clock-likeness, calculated using the variance of root-to-tip distances; 9) level of saturation, estimated as one minus the regression slope of patristic distances on *p*-distances (Nosenko, et al. 2013); 10) compositional heterogeneity, measured by the relative composition frequency variability (RCFV; Nesnidal, et al. 2010); 11) average BS support; 12) Robinson-Foulds (RF) similarity to the species tree supported by each study (Robinson and Foulds 1981); two estimates of evolutionary rates, including 13) the total tree length divided by the number of terminals (Telford, et al. 2014) and 14) the harmonic mean of site rates; and 15) the area under the penalized PI profile (iPIpen). For this last one, site rates were used to estimate a PI profile with *PhyInformR* (Dornburg, et al. 2016) for the entire time spanned between root and tips. To account for the accumulation of phylogenetic noise, informativeness values for times older than that of the peak were penalized following the method described in Bellot, et al. (2020). The area under the curve was estimated using spline interpolation with the package *MESS* (Ekstrom 2020). All properties were measured at the level of genes. Metrics were defined such that positive attributes (such as RF similarity) should be maximized, while negative attributes (such as level of saturation) should be minimized.

Across all datasets, proxies for phylogenetic signal (average BS, RF similarity and iPIpen) correlate most strongly with the length, rate of evolution and proportion of variable sites of loci, increasing with all of these (Fig. S1). Other properties previously suggested as strong predictors, such as clock-likeness and compositional heterogeneity (Doyle, et al. 2015; Shen, et al. 2016; Kuang, et al. 2018; Vankan, et al. 2020; Evangelista, et al. 2021), show less predictable relationships that can range from strongly positive to strongly negative (Fig. S1). Some variables (e.g., saturation, treeness) have stronger effects on some proxies than others, which further complicates extracting meaningful patterns. More importantly perhaps, 97.1% of all pair-wise correlations among the 15 properties are significant across more than half of the datasets (including those between signal proxies and all predictors; Fig. S2).

In order to explore whether gene properties share common patterns of covariance across datasets, I followed the approach of Mongiardino Koch and Thompson (2020), focusing on a subset of seven variables: two proxies for signal (average BS and RF similarity), four sources of bias (average pair-wise patristic distance, level of saturation, root-to-tip variance and compositional heterogeneity), and the proportion of variable sites. A principal component analysis (PCA) of these datasets resulted in two major axes explaining an average of 51.7 and 24.5% of total variance. Hierarchical and *k*-means clustering of the loadings of these first two principal components (PCs) support the hypothesis that these axes are capturing similar aspects of molecular evolution across datasets (Figs. 1 and S3). Both techniques resulted in a split of PCs into two main groups: one that includes PCs along which all properties increase/decrease (a pattern generally captured by PC 1), and another group of PCs along which sources of bias change in the opposite direction than proxies for signal (a pattern generally retrieved as PC 2). Two datasets (Hexapoda and Phasmatodea) have PCs whose groupings are reversed relative to others.

**Figure 1:**
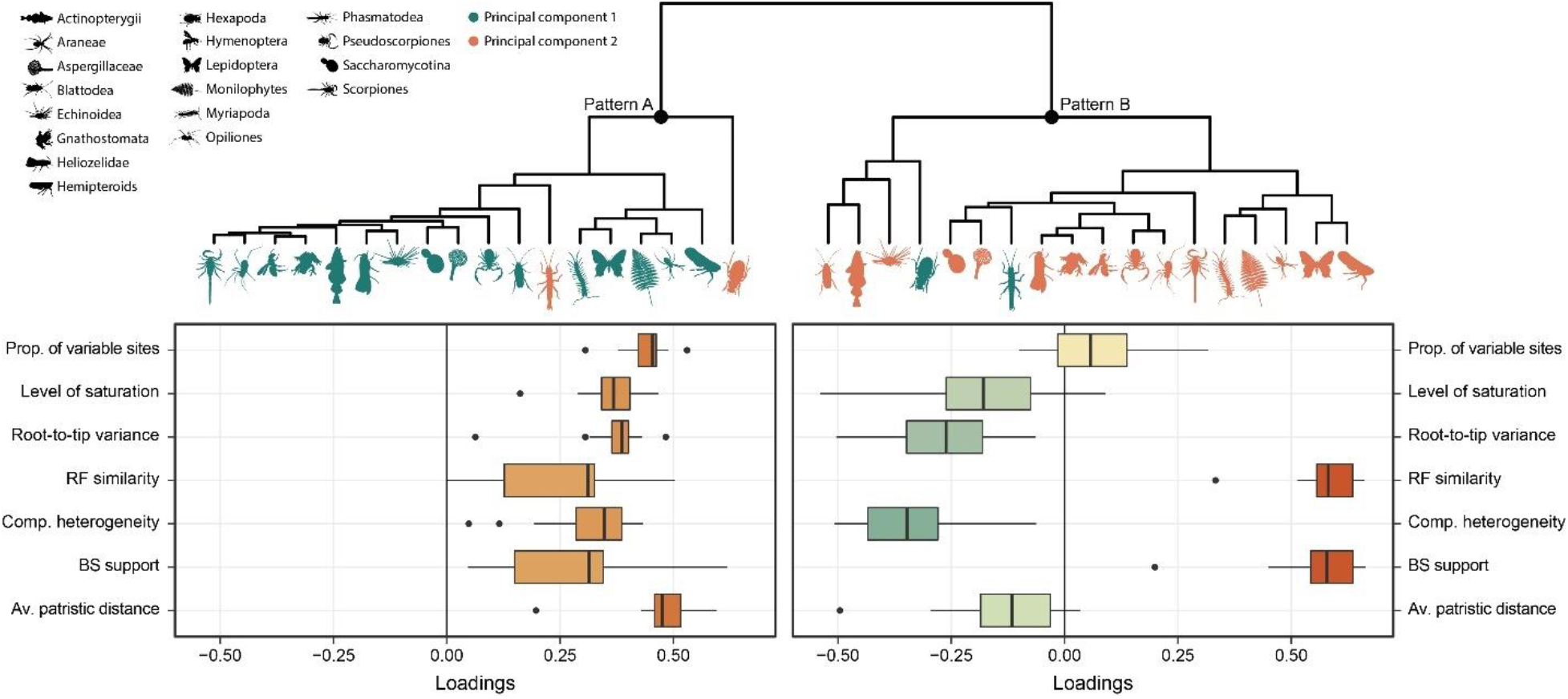
Gene properties covary in predictable ways, revealing underlying patterns of evolution that are shared by all phylogenomic datasets. The dendrogram shows that the eigenvectors of PC axes can be clustered into two major groups, labelled as patterns A and B. While pattern A is generally captured by PC 1 (green icons) and pattern B by PC 2 (orange icons), the hexapod and phasmatodean datasets are inverted. The histograms on the bottom she the distribution of loadings across variables. Results using *k*-means clustering are shown in Fig. S3.

To understand what underlying factors could be generating these patterns, the scores of loci along both PCs were correlated with estimates of evolutionary rates. This analysis confirmed that the variability generally captured along PC 1 reflects differences in rates of evolution (Fig. 2). On the other hand, PC 2 constitutes a dimension that is largely uncorrelated with evolutionary rates, but that often shows a more or less conspicuous peak at intermediate rates. Once again, the hexapod and phasmatodean datasets deviate from these patterns by exhibiting the lowest levels of correlation between rates and PC 1, as well as the highest level of correlation between rates and PC 2 (in absolute terms). These results are insensitive to the choice of an alternative, tree-based method to estimate evolutionary rates (Fig. S4).

**Figure 2:**
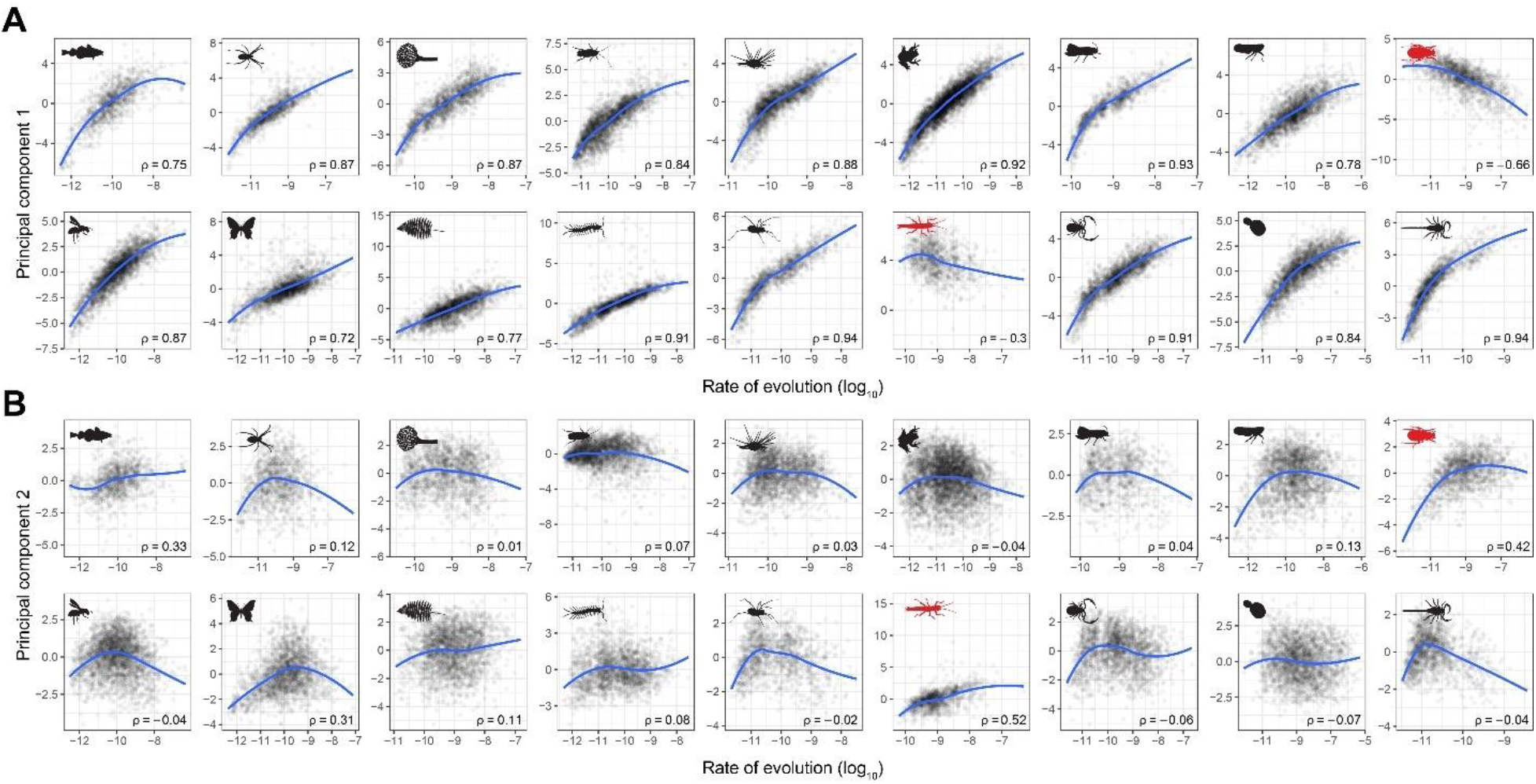
Rate of evolution is the primary factor driving differences in gene properties. Scores of loci along principal components 1 (A) and 2 (B) were correlated against the log-transformed harmonic means of site rates. Blue lines correspond to LOESS regressions, and Spearman’s rank correlation coefficients (ρ) are shown in each plot. Clade icons are as in Fig. 1; the deviating hexapod and phasmatodean datasets are highlighted in red. Results using a tree-based estimate of evolutionary rates are shown in Fig. S4.

The phylogenetic behavior of loci selected by both PC axes was then compared against other common subsampling strategies. For this, phylogenomic datasets were sorted according to a number of criteria and reduced to sizes of both 50 and 250 loci, selecting those that scored the highest or the lowest, depending on the strategy. A total of 23 subsampled matrices of both sizes were built from each dataset. These included matrices that maximized gene length, occupancy, proportion of variable sites, average BS, RF similarity, iPIpen and treeness, as well as matrices that minimized saturation, compositional heterogeneity and root-to-tip variance. Datasets were also built from the fastest and slowest evolving loci, those showing intermediate rates (i.e., those whose rates were closest to the median rate of the entire dataset), as well as those that scored highest and lowest along PC axes 1 and 2. Sorting was also done with SortaDate (Smith, et al. 2018), a common pipeline for phylogenomic subsampling based on three gene properties. However, this method ordered loci in ways that were nearly identical to those achieved by using just one variable, whichever was selected as the first sorting step (see Fig. S5). Since all three variables were already being assessed, this method was not employed. Finally, five datasets were generated by sampling genes at random.

Phylogenetic inference using subsampled datasets was performed using IQ-TREE 1.6.3 (Nguyen, et al. 2014) under the LG+F+G model, and node support was estimated using 1,000 replicates of ultrafast bootstrap (UFBoot; Hoang, et al. 2017). Characterizing the performance of these datasets is complicated by the fact that the underlying phylogenies are unknown (in fact some of the trees used here have already been challenged to some degree; see Meusemann, et al. 2020; Szucsich, et al. 2020; Tihelka, et al. 2020). While large phylogenomic datasets generally produce fully resolved and supported topologies, model violations can favor incorrect trees (Delsuc, et al. 2005; Kapli, et al. 2021). While this necessarily means that topologies supported by full phylogenomic datasets are only imperfect proxies with which to evaluate phylogenetic accuracy, it is also true that the proportion of nodes sensitive to model choice in any given analysis is small. Optimal subsampled datasets should be able to recapitulate this general tree structure, although not necessarily every detail; in other words, high topological similarity should still be favored, although the highest value does not guarantee the best results. At the same time, genes differ in their levels of phylogenetic signal, and an adequate subsampling scheme should be able to recover genes with above-average performance. Considering this, subsampling schemes were ranked in descending order of RF similarity to the tree found by the original studies, breaking ties using the average UFBoot values. The values for the five replicates of random subsampling were averaged to obtain a single estimate of their performance. Subsampling strategies ranking systematically better than random-chosen loci were considered valid. Given difficulties establishing the identity of PC axes for Hexapoda and Phasmatodea, the results of these datasets were not included with the rest and are shown separately in Fig. S6.

When subsampling to 250 loci, only five methods outperformed randomly-chosen loci across more than half of the datasets (Fig. 3A). These include matrices designed to maximize RF similarity, average BS, occupancy and length, as well as those with loci that rank highest along PC 2. Two additional approaches—iPIpen and intermediate rates—have median ranks above that of randomly chosen loci, although ranking below more often than not. Of these, RF similarity and PC 2 (high) are the most consistent (i.e., have the lowest variance); other approaches behave well on average, but can occasionally perform poorly. As expected, differences in performance between strategies are even larger when subsampling to 50 loci (Fig. S7); however, the same set of methods are favored, with the further addition of loci with the highest proportions of variable sites. Very common approaches, including rate-based subsampling (saving the marginally good behavior shown by loci with intermediate rates) and the direct minimization of systematic biases (including saturation and among-lineage compositional and rate heterogeneities), perform systematically worse than randomly-chosen loci at both subsampling levels (Figs. 3A and S6).

**Figure 3:**
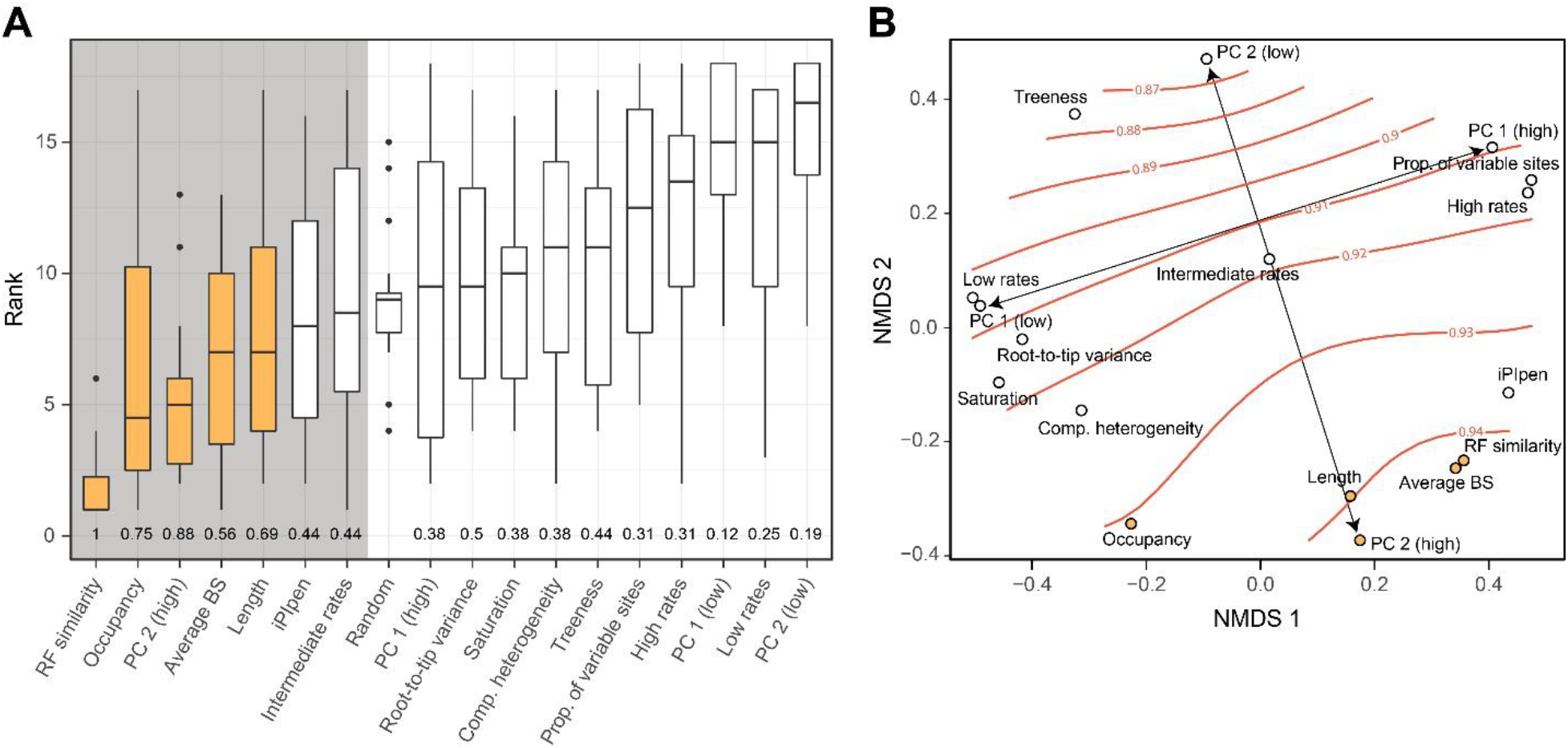
Comparison of the performance of alternative subsampling strategies. **A.** Distribution of ranks attained by different strategies (lower ranks represent better results). Two criteria for selecting adequate strategies are highlighted: those whose median ranks are lower than randomly chosen loci (grey background), and those that outperform these in more than half of the datasets (yellow bars). The proportion of times a given strategy ranks better than random loci is shown at the bottom. Results correspond to matrices of 250 loci; those for 50 loci are shown in Fig. S7. **B.** Non-metric multidimensional scaling (NMDS) of pair-wise distances between strategies, representing the average frequency with which they share loci (smaller distances represent higher probabilities of targeting the same loci). Average RF similarity (orange lines) is overlayed as a smooth surface. Principal component (PC) 2 defines an axis that traverses the RF similarity gradient, while PC 1 (and other rate proxies) sample genes along a perpendicular axis that follows an isocline.

To further explore these patterns, I calculated the fraction of shared loci between matrices built using different subsampling strategies. This value was turned into a pair-wise distance metric and averaged across datasets, producing an estimate of the expected frequency with which strategies select the same genes. Non-metric multidimensional scaling was used to project these distances into a two-dimensional space on which the average topological similarity was overlain (Fig. 3B). In line with previous results (Figs. 1, 2), this confirms that: 1) PCs built from the gene property datasets represent axes of evolutionary rate and phylogenetic usefulness; 2) rate and usefulness are perpendicular axes, such that rate-based subsampling does not optimize usefulness; and 3) directly minimizing sources of bias performs poorly because it has the unintended consequence of targeting slow-evolving loci that are largely uninformative.

## Discussion

Quantifying and predicting which loci contribute toward recovering correct topologies has become central to phylogenomic inference (Salichos and Rokas 2013; Doyle, et al. 2015; Edwards 2016; Shen, et al. 2016; Arcila, et al. 2017; Brown and Thomson 2017; Shen, et al. 2017; Molloy and Warnow 2018; Smith, et al. 2018; Dornburg, et al. 2019). This step can be used to explore phylogenetic conflicts, test specific hypotheses of relationships, measure the impact of different sources of bias, and allow for a better modelling of evolutionary processes. For the many phylogenetic questions that still remain unanswered, the preferred topology can entirely depend on assessments of the phylogenetic information contained within different loci (e.g., Simon, et al. 2018; Lozano-Fernandez, et al. 2019; Marlétaz, et al. 2019; Smith, et al. 2020). This has led to a plethora of recommendations on what constitutes a reliable gene and which proxies can be used to enrich datasets in them. Many of these were supported by searching for strong predictors of the topological distance to a preferred topology (Doyle, et al. 2015; Burbrink, et al. 2020; Vankan, et al. 2020). However, extracting the individual effects of potential predictors is complicated by the pervasive levels of correlation that these exhibit (Shen, et al. 2016; Kocot, et al. 2017; Mongiardino Koch and Thompson 2020). Subsampling based on any individual property in the presence of such strong correlations can also have unintended effects: for example, increasing occupancy can reduce overall levels of phylogenetic signal, and targeting longer genes can increase compositional heterogeneity (Figs. S1, S2).

Instead of focusing on correlating pairs of variables, I propose that a better understanding of the information content of loci can be gained by searching for regularities in the patterns of covariance between multiple properties, and exploring the underlying factors that might produce them. Across a sample of 18 diverse phylogenomic datasets, I find that most of the variability captured across multiple gene properties happens along two major axes. These axes show remarkably similar patterns of covariance that can be readily interpreted as representing differences in evolutionary rate and phylogenetic usefulness (Figs. 1, 2, S4, S6). In the case of the latter, highly useful loci exhibit a consistent set of properties that include not only high values of node support and topological similarity, but also low levels of saturation and reduced compositional and rate heterogeneities (i.e., simultaneously high signal and low biases). They also seem not to be among the fastest or slowest evolving genes, implying the existence of an optimal rate as predicted by theory (Yang 1998; Townsend 2007; Susko and Roger 2012; Klopfstein, et al. 2017; Dornburg, et al. 2019). Datasets with high levels of rate variation have reduced variation in phylogenetic usefulness and vice versa (Fig. S8), which is also expected if usefulness peaks at a particular (optimal) rate.

Many common subsampling strategies are justified in either phylogenetic theory or in the aforementioned correlation with measures of topological distance at the gene level. However, the behavior of multi-locus subsampled datasets obtained by filtering genes based on such correlates has been seldom explored. Phylogenetically useful loci should also possess other properties besides low topological distances to a target tree, such as displaying a minimum of non-phylogenetic signals that can provide hidden support for an incorrect topology when loci are concatenated (Gatesy and Springer 2014). When the performance of subsampling strategies is evaluated, it becomes clear that many common approaches do not perform well. Such is the case of rate-based subsampling: matrices composed of the slowest or fastest evolving loci are among the worst that can be generated from phylogenomic datasets (Fig. 3, S6). Even targeting loci with intermediate rates, or those that evolve at a pace that should maximize phylogenetic informativeness, does not drastically improve results relative to selecting loci at random (although iPIpen does succeed when subsampling to very small sizes; Fig. S7). Different lines of evidence show that this inefficacy is a consequence of evolutionary rate being a dimension that is perpendicular to phylogenetic usefulness (Figs. 1, 3B). At first glance, this might seem to conflict with the existence of optimal rates for inference, but peaks in usefulness are evident in Figures 2 and S4.

Another explanation could be that a direct link between rates and usefulness only exists at the level of sites (Dornburg, et al. 2019), as entirely different distributions of site rates can potentially average to identical gene rates. This not only implies that gene rates should be avoided for subsampling, but they might even constitute abstractions with weak ties to evolutionary processes. The results presented here confirm that gene rates are not a useful subsampling approach, but they also show that they do capture relevant differences in evolutionary history. Multiple proxies for gene rates seem to converge on similar values, and genes with comparable rates share many common features, defining the major axis of variance in gene properties across most datasets. The problem does not seem to lie in gene rates being inappropriate, but rather that they constitute just one of several criteria that a phylogenetically useful locus should possess. Loci evolving at optimal gene rates exhibit large variabilities in usefulness (Fig. S9), which makes rate-based subsampling inefficient even when optimal gene rates can be discovered. While this might be caused by differences in the underlying distributions of site rates, it likely also reflects compositional biases and rate heterogeneities that are not accommodated by approaches based on rates or informativeness (Dornburg, et al. 2019).

Another common method to reduce the size of phylogenomic datasets is to discard loci that seem most affected by potential sources of bias (Nesnidal, et al. 2010; Borowiec, et al. 2015; Whelan, et al. 2015; Kocot, et al. 2017; Mongiardino Koch, et al. 2018; Marlétaz, et al. 2019), including high levels of saturation and heterogeneities in both composition and evolutionary rates. However, selecting the loci least affected by these issues does not result in phylogenetically accurate datasets (Fig. 3A). These results are in strong conflict with many previous analyses that supported the use of clock-like, unsaturated and compositionally homogenous genes (Doyle, et al. 2015; Kuang, et al. 2018; Lozano-Fernandez, et al. 2019; Vankan, et al. 2020; Evangelista, et al. 2021). While all three of these properties clearly represent severe issues for phylogenetic inference (Delsuc, et al. 2005; Kapli, et al. 2021), directly minimizing them enriches the dataset in conserved and slow-evolving loci that do not contain enough phylogenetic information (Fig. 3B). This unintended consequence highlights the fact that selecting genes based on any individual attribute can produce strong and undesired shifts in the distributions of other variables. Clock-like genes are also routinely favored for estimating divergence times (Smith, et al. 2018; Carruthers, et al. 2020); it is therefore important to note that sampling the most clock-like genes can deplete phylogenetic signal and bias rate estimates.

Only five approaches are found to systematically outperform random loci selection at both levels of subsampling (Figs. 3, S6). These include two proxies for phylogenetic signal (RF similarity and average BS), two measures of amount of information (alignment length and occupancy), and the phylogenetic usefulness axis obtained using PCA. The finding that maximizing RF similarity is consistently recovered as the best approach was expected, as the ranking of strategies is to a large degree also determined by this metric. This circularity complicates an objective evaluation of this approach, which would require simulations under a known topology (to some degree, this is true for other conclusions drawn here). However, maximizing average BS support, a different proxy for signal that does not suffer from this problem, results in the sampling of a very similar set of loci (Fig. 3B), providing indirect evidence of the suitability of subsampling based on topological similarity. At the same time, given that sampling of genes selected for their RF similarity recovers the topologies most similar to those of targeted trees, this strategy provides an effective way of replicating results with smaller datasets, but should not be interpreted as a test of phylogenetic results. While longer genes were previously found to recover better topologies (Aguileta, et al. 2008; Shen, et al. 2016; Brown and Thomson 2017), occupancy had been considered less of a concern for datasets composed of hundreds of loci (Philippe, et al. 2004; Streicher, et al. 2016; Molloy and Warnow 2018). Results shown here suggest that maximizing both of these are among the best-performing subsampling strategies on average, but also exhibit a relatively inconsistent behavior, occasionally ranking among the worst. Their use should be accompanied by some assessment of how they are impacting overall levels of signal.

Finally, maximizing phylogenetic usefulness through the use of PCA provides a direct way to optimize levels of phylogenetic signal while also controlling for sources of bias. This is done simultaneously and without the need to arbitrarily order variables or establish thresholds. By drawing information from multiple properties, the approach is able to discover patterns that are unique to each dataset, weighting factors in proportion to their relative contributions. This also provides a useful avenue for filtering outlier genes, as shown in the Methods section and Figure 4. For all but two of the datasets analyzed, the interpretation of the second PC dimension as a usefulness axis was straightforward; for the remaining ones (Phasmatodea and Hexapoda), a more careful study revealed usefulness was captured along PC 1 (Figs. S6). In the specific case of the hexapod dataset, both PC axes seemed to correlate relatively strongly with rate estimates (Fig. 2), which is consistent with the idea that resolving the phylogeny of ancient clades requires highly conserved, slow-evolving genes. Taken to an extreme, this could potentially induce the collapse of rate and usefulness into a single dimension, at which point the method here described would become impractical. Therefore, this approach may not be universally applicable, and might not help resolve phylogenies outside the range of conditions explored, including clades that are older, evolve faster, or contain recalcitrant nodes. Even so, it is likely that progress in our understanding of contentious relationships that have defied resolution will happen as we improve our ability to decode the evolutionary processes ingrained along the different axes that describe the information content of loci.

**Figure 4:**
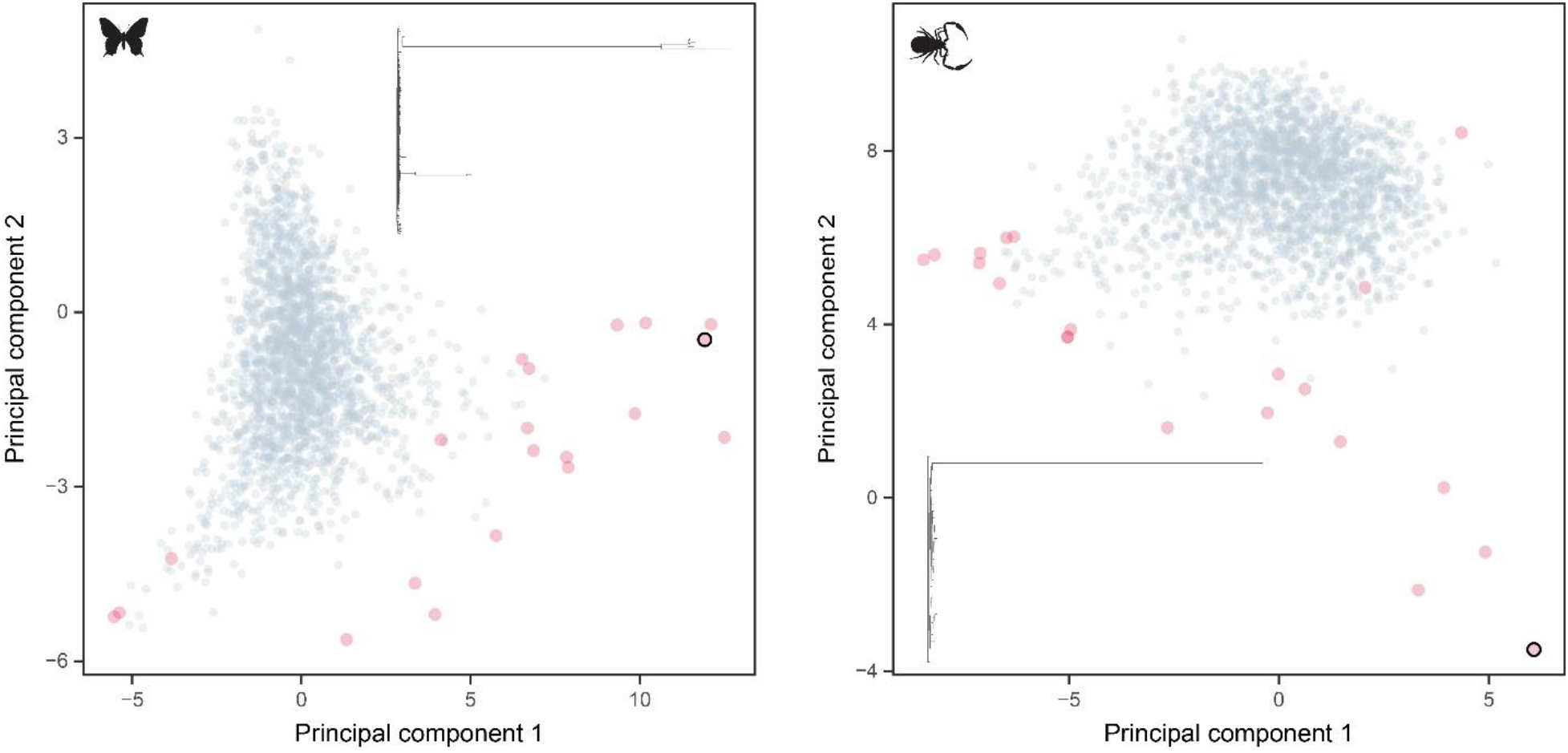
Detection of outlier genes using multiple gene properties in two exemplary datasets, Lepidoptera (left) and Pseudoscorpiones (right). Plots shows the principal component axes built from the entire datasets, with the genes considered outliers shown in red. The topology of the largest outlier (highlighted with a black border) is plotted.

## Methods

Datasets chosen for this study had to fulfill a number of criteria. First, I only used datasets built from full genomes and/or transcriptomes, as these are likely to exhibit a wider range of values across different properties—such as rates—than datasets built using methods of targeted enrichment (e.g., ultraconserved elements, anchored hybrid enrichment). For standardization, all datasets were coded as amino acids, although the methods employed are applicable to other data types. Studies also had to infer a time-calibrated topology, establishing a timescale of diversification that could be used to estimate rates of evolution in number of substitutions per unit of time. These topologies were inferred and calibrated using entirely different methodologies, but represent in every case the best estimate of relationships as supported by the authors. Finally, datasets with notoriously contentious relationships, such as chelicerates (Sharma, et al. 2014) and metazoans (King and Rokas 2017), were avoided. The 18 datasets sampled (Table 1) were only modified by filtering loci with values of occupancy below 50%.

Gene trees were inferred using ParGenes v. 1.0.1 (Morel, et al. 2018) which automated model selection with ModelTest-NG (Darriba, et al. 2020) and phylogenetic inference with RAxML-NG (Kozlov, et al. 2019) for each multiple sequence alignment. The optimal model was considered to be the one minimizing the Bayesian Information Criterion; support values were estimated with 100 replicates of non-parametric bootstrap (BS). Rates of evolution for all sites in each datasets were estimated using the empirical Bayes method implemented in Rate4Site (Mayrose, et al. 2004) using the time-calibrated tree pruned to include only terminals present in each locus. Given that outgroups often represent poorly-sampled clades that can be distantly related to the ingroup (e.g., in the case of Echinoidea extending the age of the tree root by 200 million years; Mongiardino Koch and Thompson 2020), and thus have a strong effect on estimated rates, they were removed from both trees and alignments. Branch length optimization was disabled and all other options were left as default. For some loci, the inference of gene trees or the estimation of site rates failed; these loci were dropped from further analyses, resulting in the final numbers shown in Table 1.

A group of 15 properties were calculated for each locus in R using custom scripts (see Results). Scripts relied on functions from packages *adephylo* (Jombart and Dray 2010), *ape* (Paradis and Schliep 2018), *MESS* (Ekstrom 2020), *phangorn* (Schliep 2011), *PhyInformR* (Dornburg, et al. 2016), *phytools* (Revell 2012) and the *tidyverse* (Wickham 2017). As with site rates, outgroups were removed before estimating these. Correlations were visualized using package *corrplot* (Wei and Simko 2017), and *P* values were corrected using Benjamini and Hochberg (1995) correction for multiple comparisons.

Following Mongiardino Koch and Thompson (2020), a subset of seven gene properties were subject to principal component analysis (PCA). Among these are two widely employed proxies for phylogenetic signal: the Robinson-Foulds (RF) similarity to the species tree (i.e., the complement of the RF distance; Robinson and Foulds 1981), generally taken to be an estimate of topological accuracy, and the average BS support (Salichos and Rokas 2013; Doyle, et al. 2015; Shen, et al. 2016; Vankan, et al. 2020). Four other variables are known to induce systematic errors in tree reconstruction (Delsuc, et al. 2005; Nesnidal, et al. 2010; Nosenko, et al. 2013; Struck 2014; Kocot, et al. 2017; Kapli, et al. 2021): the variance of root-to-tip distances (i.e., the degree of deviation from a strict clock-like behavior), the average pair-wise patristic distance between terminals (indicative of susceptibility to long-branch attraction), the level of saturation (estimated as one minus the regression slope of patristic distances on *p*-distances), and the compositional heterogeneity (measured by the relative composition frequency variability; RCFV). The last variable included was the proportion of variable sites, a metric generally interpreted to represent information content (Aguileta, et al. 2008; Mclean, et al. 2019), and that is strongly correlated with estimates of rates and tree length in the datasets employed (Fig. S2). All of these properties have been used individually for phylogenomic subsampling. This approach suffers from some degree of circularity given the use of topological similarity in the selection of genes, but this should bias results minimally as this is just one of the several attributes employed. In case this represents a concern, uncertain nodes can be collapsed in the tree used to measure topological distances; taken further this would converge on the approach used by Philippe, et al. (2019) to focus only on the recovery of a handful of uncontroversial monophyletic groups. A few different sets of variables were explored, as well as alternative metrics for some of them (such as different tree distances); these changes did not improve the proportion of variance captured by the first two principal components (PCs) and were not further explored. It should be noted however that a thorough optimization of the variables included was not performed, and this is likely to have some effect on results.

PCA is susceptible to outlier data points (i.e., observations that strongly deviate from the general structure of correlation between variables), as these contribute a large fraction of total variance and can attract the first components. Although this can be seen as a limitation of the method, it also provides an opportunity to detect and filter out outlier genes. These can arise from both analytical and biological processes (e.g., errors in orthology inference or alignment, strong selective pressures, etc.), and have a strong impact on tree reconstruction (Brown and Thomson 2017; Shen, et al. 2017; Walker, et al. 2018). To remove outlier genes, I measured the Mahalanobis distance of all observations to the origin of the PC space (employing all seven dimensions), and removed the top 1% with the greatest distances (Fig. 4).

These represent sequences with highly unlikely combinations of gene properties given the structure of correlation of the entire dataset. PCA was then repeated on the remaining observations. Compared to other methods devised to remove outlier data from phylogenomic datasets (e.g., de Vienne, et al. 2012; Mai and Mirarab 2018), this approach benefits from not only considering tree topology, but doing so alongside other gene properties. The removal of outlier genes not only helps correctly identify the major axes of variance among ‘regular’ observations (i.e., ensures that PCs capture true differences in rate and usefulness), but also provides an extra step of sanitation, likely to be especially important before datasets are reduced in size. Future work would likely benefit from a more sophisticated approach to outlier detection, such as is offered by robust PCA methods (Todorov and Filzmoser 2009).

Both hierarchical and *k*-means clustering were used to discover groupings of similar PC axes that could potentially represent similar underlying factors. Given that PC orientation is arbitrary, clustering was done using eigenvectors as well as their opposites (Fig. 1 has the mirrored half of the dendrogram removed). Hierarchical clustering was performed using Euclidean distances and complete linkage (Fig. 1); *k*-means clustering used 10,000 random starting configurations (Fig. S3). The identity of these axes was first established by correlating the scores of the first two PCs against different estimates of gene-wise evolutionary rates: the total tree length divided by the number of terminals (Telford, et al. 2014; Howard, et al. 2020), and the harmonic mean of site rates. For all datasets except Hexapoda and Phasmatodea, the Spearman rank correlation coefficients (ρ) between both estimates of rate and PC 1 were larger than 0.7 and more than twice the values of ρ between rate estimates and PC 2 (Figs. 2, S4). This was taken to represent strong evidence that PC 1 was (in general) capturing rate variation. Correlations between PC 1 and tree-based rates were much higher (average ρ = 0.94) than between PC 1 and sequence-based rates (average ρ = 0.86). This seems to confirm that averaged site rates are an inaccurate proxy for gene-wise evolutionary rates (Dornburg, et al. 2019). The relationship between gene rates and phylogenetic usefulness (Fig. S9) was also studies by binning loci into 25 categories based on their rates, and calculating the mean and variance of usefulness (i.e., PC 2 scores) within each. A linear regression between these two metrics was assessed after excluding outliers, identified as those whose residuals were significantly larger than expected using a chi-square test in package *outliers* (Komsta 2011).

Phylogenomic datasets were sorted based on 13 different properties (gene length, occupancy, proportion of variable sites, average BS, RF similarity, iPIpen, treeness, saturation, RCFV, root-to-tip variance, sequence-based evolutionary rate, and PCs 1 and 2) and subsampled to sizes of 50 and 250. These numbers were chosen because they represent common data sizes used for computationally intensive methods such as total-evidence dating (Lee 2016; Brennan, et al. 2020; Mongiardino Koch and Thompson 2020) and inference under complex site heterogenous models (Ballesteros, et al. 2019; Marlétaz, et al. 2019), respectively. Subsampled datasets were composed of either the highest or lowest scoring loci, depending on the variable used for sorting. In the case of rates and PC axes, both the highest and lowest scoring loci were used. An extra subsampling strategy targeting intermediate rates (defined as those loci with sequence-based rates closest to the median value for the entire dataset) was also used. Five extra matrices were built by selecting loci at random, for a total of 23 matrices per phylogenomic dataset and subsampling size. Tree inference was performed in IQ-TREE 1.6.3 (Nguyen, et al. 2014) under the LG+F+G model, and 1,000 replicates of ultrafast bootstrap (UFBoot; Hoang, et al. 2017) were used to estimate node support values.

The performance of subsampling strategies was evaluated using two metrics: the RF similarity to the tree supported by the original studies (i.e., the same used to estimate topological similarity for individual loci), and the average UFBoot support. The values obtained for the five replicates of randomly sampled loci were averaged. Subsampling strategies were then ranked based on RF similarity scores with ties broken using average support values, such that strategies that result in more accurate and well-supported trees receive lower ranks. Two criteria were used to establish which subsampling approaches are useful: 1) strategies that attain a median rank that is lower than that of randomly sampled data across datasets; and more strictly, 2) strategies that attain a lower rank than randomly sampled data for more than half of datasets (Figs. 3, S7). Given the non-standard behavior of the hexapod and phasmatodean datasets, results from these were not combined with those of other datasets, and are reported separately in Figure S6. It should be noted that subsampling was always performed by selecting entire genes, and that results for some strategies might differ from those obtained by selecting sites (e.g., when using rates). Retaining the gene structure of the datasets is not only necessary for some types of phylogenetic inference such as summary coalescent methods, but also provides access to a much larger pool of properties, including all of those estimated on gene trees. A focus on loci can also help discover outlier data (Fig. 4), and reveal important evolutionary processes such as compositional and rate heterogeneities (or at least aid in their discovery). The relative performance of strategies was also evaluated at the level of the entire tree topology, and some of the methods used (e.g., iPIpen) might be more suitable for finding optimal loci to resolve specific nodes or time intervals.

Finally, the dissimilarities between pairs of 250-loci matrices obtained through different subsampling strategies (i.e., the proportion of loci not shared) was calculated and averaged across datasets. The resulting distance matrix was decomposed into a two-dimensional space using non-metric multidimensional scaling (NMDS). This relied on package *vegan* (Oksanen, et al. 2019) and employed 10,000 iterations from random starts. Stress was evaluated using a Shepard diagram (i.e., a plot of observed distances vs. ordination distances), and a non-metric estimate of goodness-of-fit returned an R squared value of 0.99. The averaged RF similarity across datasets was overlain onto this plot as a smooth surface, which was fitted using penalized regression splines.

## Acknowledgments

This paper benefitted from discussions with Casey W Dunn, Jesus Lozano-Fernandez, Paschalis Natsidis, Mattia Giacomelli, Alejandro Damián Serrano, Natasha Picciani, Jasmine Mah and Lauren Mellenthin. The manuscript was also improved by comments from Steve Haddock and Sophie Westacott. Jeffrey Townsend provided useful guidance regarding phylogenetic informativeness. I would like to thank Dominic Evangelista, Sabrina Simon, Liz Milla, Kevin P. Johnson, Bernhard Misof, Karen Meusemann, Akito Y. Kawahara, David Plotkin, Hui Shen, Rosa Fernández, Ligia Benavidez and Prashant Sharma for providing data files and helping match taxon names, as well as all other authors who made files available through online repositories. I would also like to thank the creators of Phylopic icons, including Gareth Monger, Jennifer Trimble, Melissa Broussard, Maxime Dahirel and Olegivvit. N.M.K. was supported by a Yale University fellowship.

## Data availability

R code to estimate gene properties, sort and subsample phylogenomic datasets will be deposited in GitHub. In the meantime, it can be obtained directly from the author.

---

**Figure S1:**
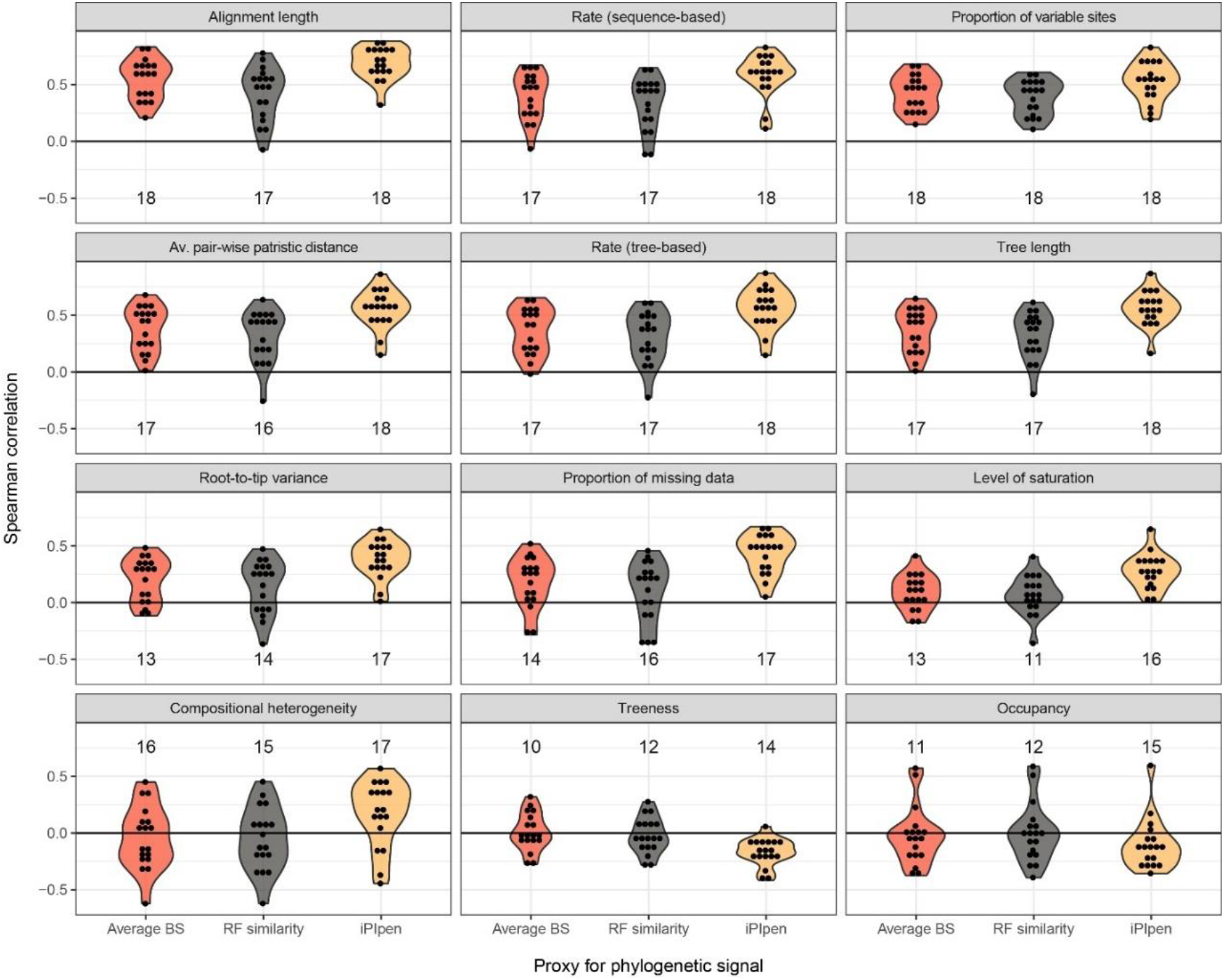
Exploring correlates of phylogenetic signal across datasets. Data shows the Spearman rank correlation coefficients between three proxies for signal (average bootstrap support, Robinson-Foulds similarity and iPIpen, the area under the penalized phylogenetic informativeness profile) and 12 other gene properties. Numbers above/below the distributions correspond to significant correlations after Benjamini & Hochberg corrections. Gene properties are ordered (top-to-bottom, left-to-right) following decreasing values of median correlation.

**Figure S2:**
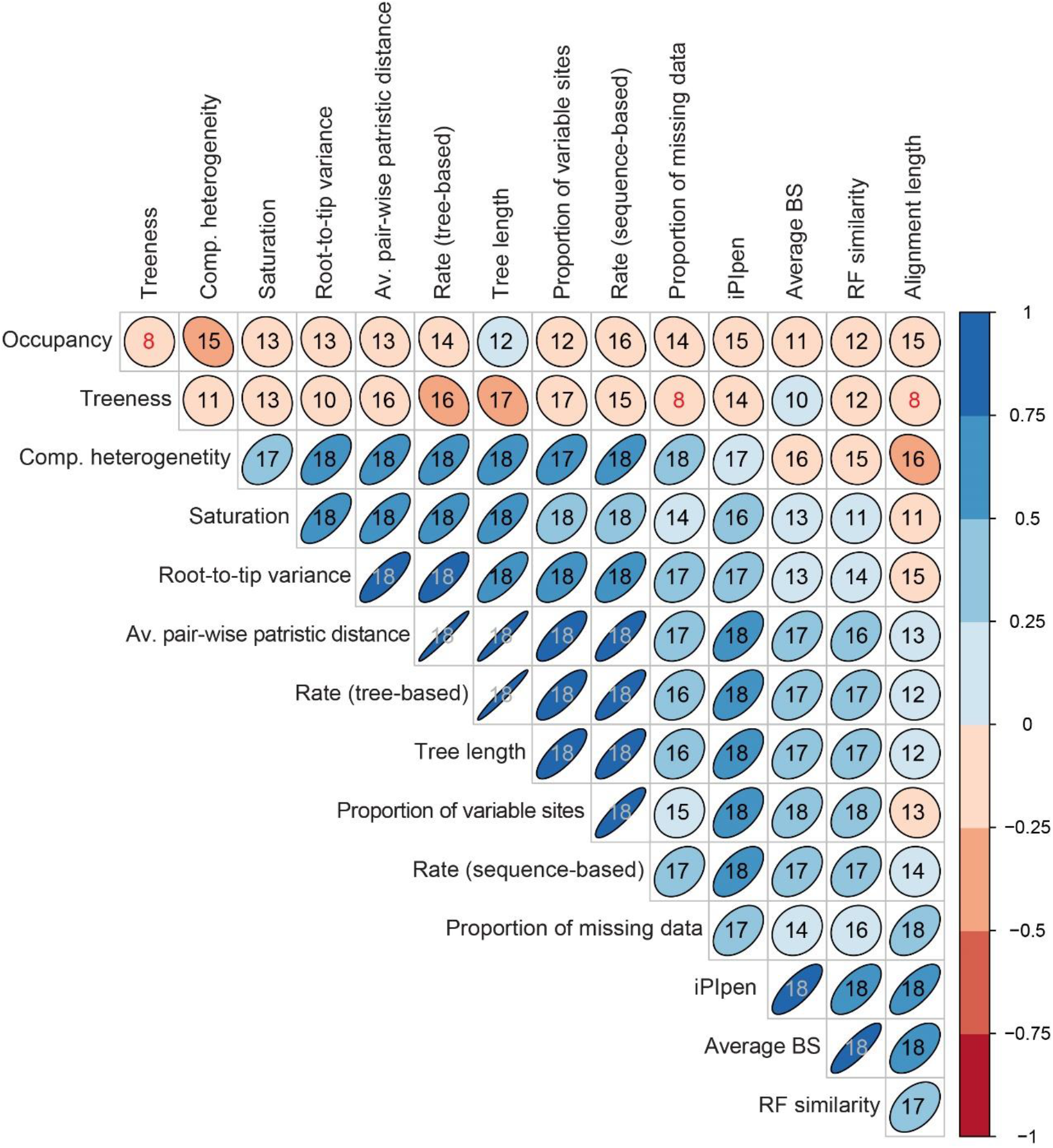
Correlation matrix for 15 gene properties. Colors and ellipse shapes correspond to average Spearman rank correlation coefficients across all 18 datasets analyzed. Number in each cell show the number of datasets for which a given correlation is significant (after Benjamini & Hochberg correction). Only 3 correlations (marked in red), all of which involve treeness, are significant in less than half of the datasets. Moderate values can arise from a true weak correlation between variables as well as an inconsistent correlation that can include both strongly positive and strongly negative values (see Fig. S1), and should therefore be interpreted with caution.

**Figure S3:**
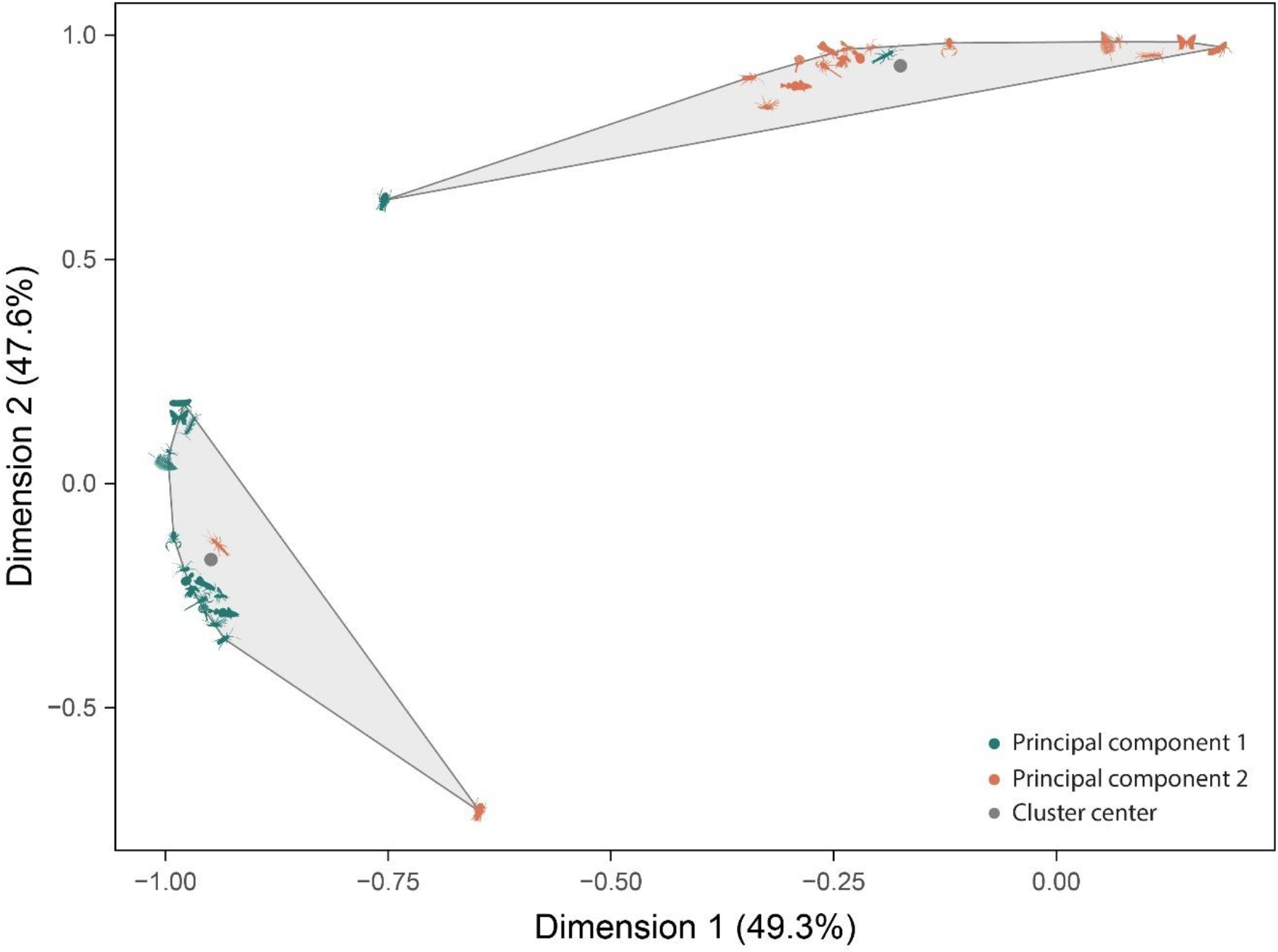
*K*-means clustering of eigenvectors. Given the results of hierarchical clustering (Fig. 1), the number of clusters was set to two. Clustering results are shown using grey convex hulls; eigenvectors for different PCs and datasets are denoted using colors (see legend) and clade icons (as in Figs. 1, 2). The eigenvectors of hexapods and phasmatodeans are clustered in an opposite pattern. PC 1 of phasmatodeans is well within the range of variability shown by the remaining PCs 2, and PC 2 is within the ranges of the remaining PCs 1. On the other hand, both hexapod eigenvectors seem to be somewhat different to those of the other datasets.

**Figure S4:**
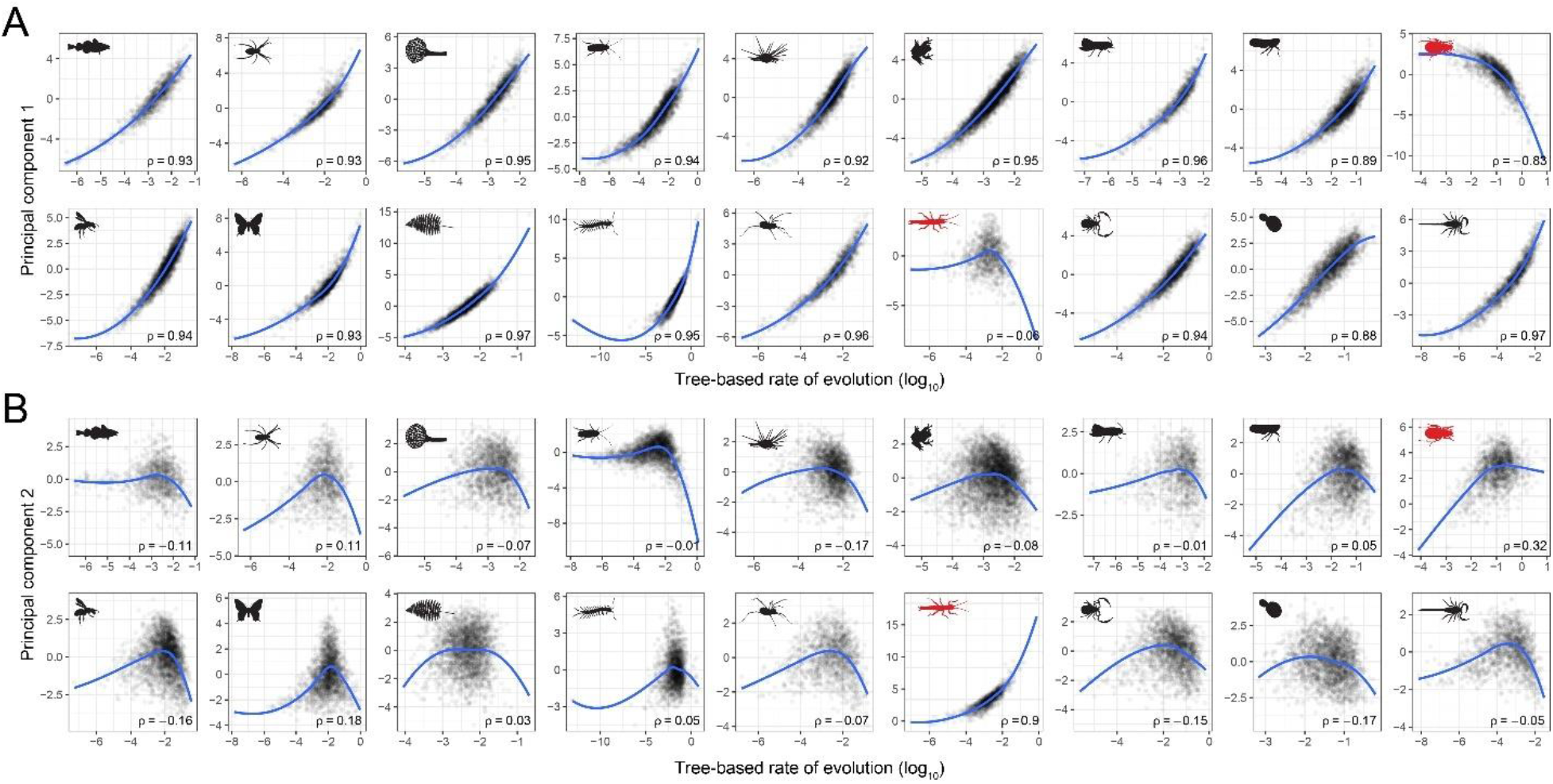
Correlation between tree-based rate of evolution and principal components 1 (A) and 2 (B). Results correspond to Fig. 2 but using an alternative, tree-based estimate of evolutionary rates. Blue lines correspond to LOESS regressions, and Spearman’s rank correlation coefficients (ρ) are shown in each plot. Clade icons are as in Figs. 1–2; the deviating hexapod and phasmatodean datasets are highlighted with red icons.

**Figure S5:**
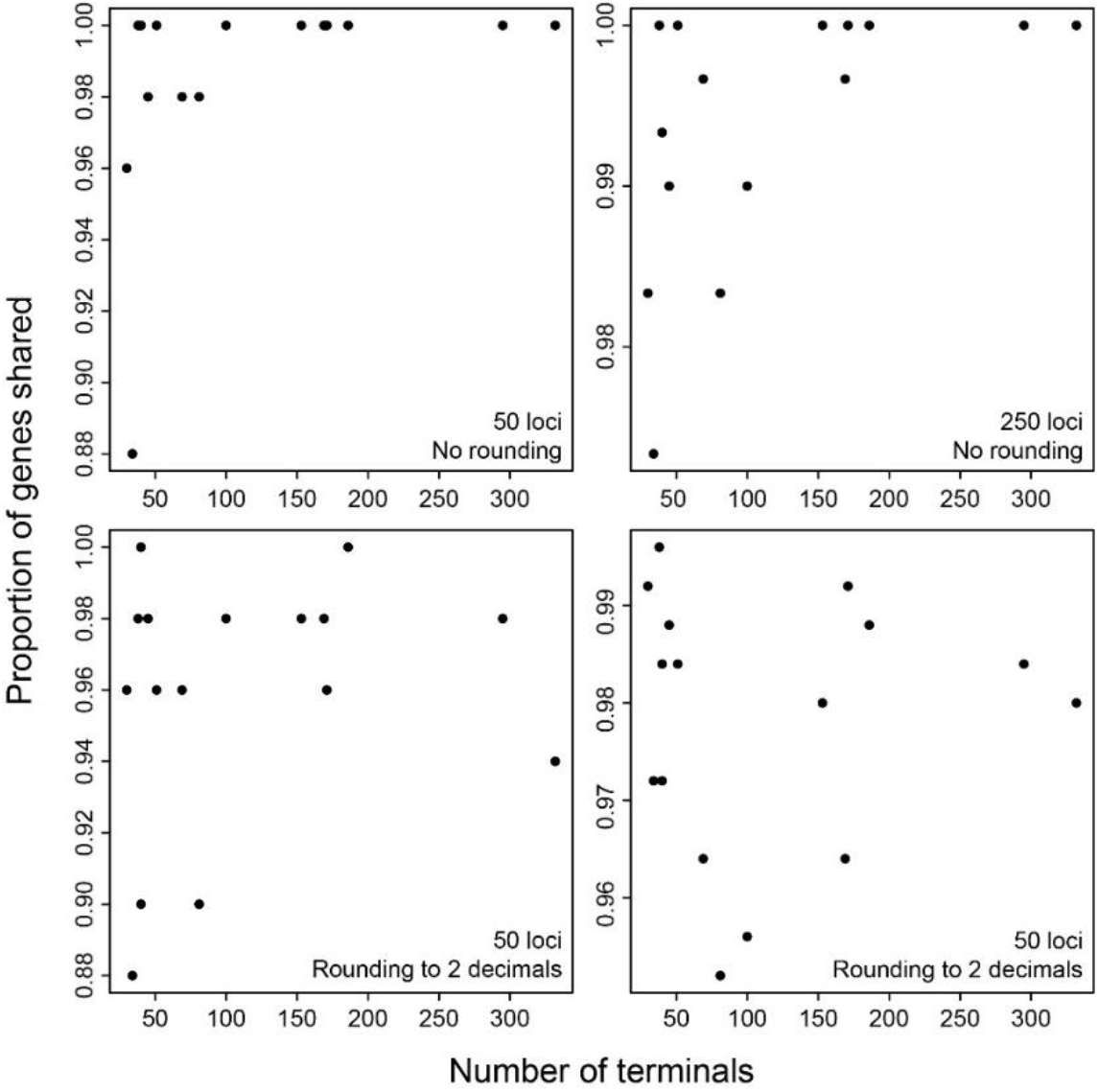
Proportion of loci shared between two strategies, SortaDate (Smith, et al. 2018) and Robinson-Foulds (RF) similarity. SortaDate uses three sequential sorting steps (topological distance, root-to-tip variance and tree length) for subsampling. Subsequent steps are only used to break ties in the previous sorting rounds; as such, it is unclear how much they contribute. Here, datasets were sorted and subsampled to 50 (left) and 250 (right) loci using all three steps of SortaDate as well as just the first one, and the proportion of genes shared between the two was calculated. The set of loci selected by SortaDate is almost entirely determined by its first step (i.e., RF similarity; top panels), with a minor impact of other variables seen only when trees have less than 100 terminals. RF similarity could potentially be rounded to force ties and increase the contribution of other variables (even though it has an intrinsic unit of scale determined by the number of terminals). However, even when rounding to 2 decimal places, the order is still almost entirely determined by the first sorting step (bottom panels). Although the order in which variables are evaluated by SortaDate can be modified, results are the same: the genes chosen are almost entirely determined by the first variable (results not shown). Note that SortaDate uses the proportion of concordant bipartitions instead of RF similarity. This does not modify these results.

**Figure S6:**
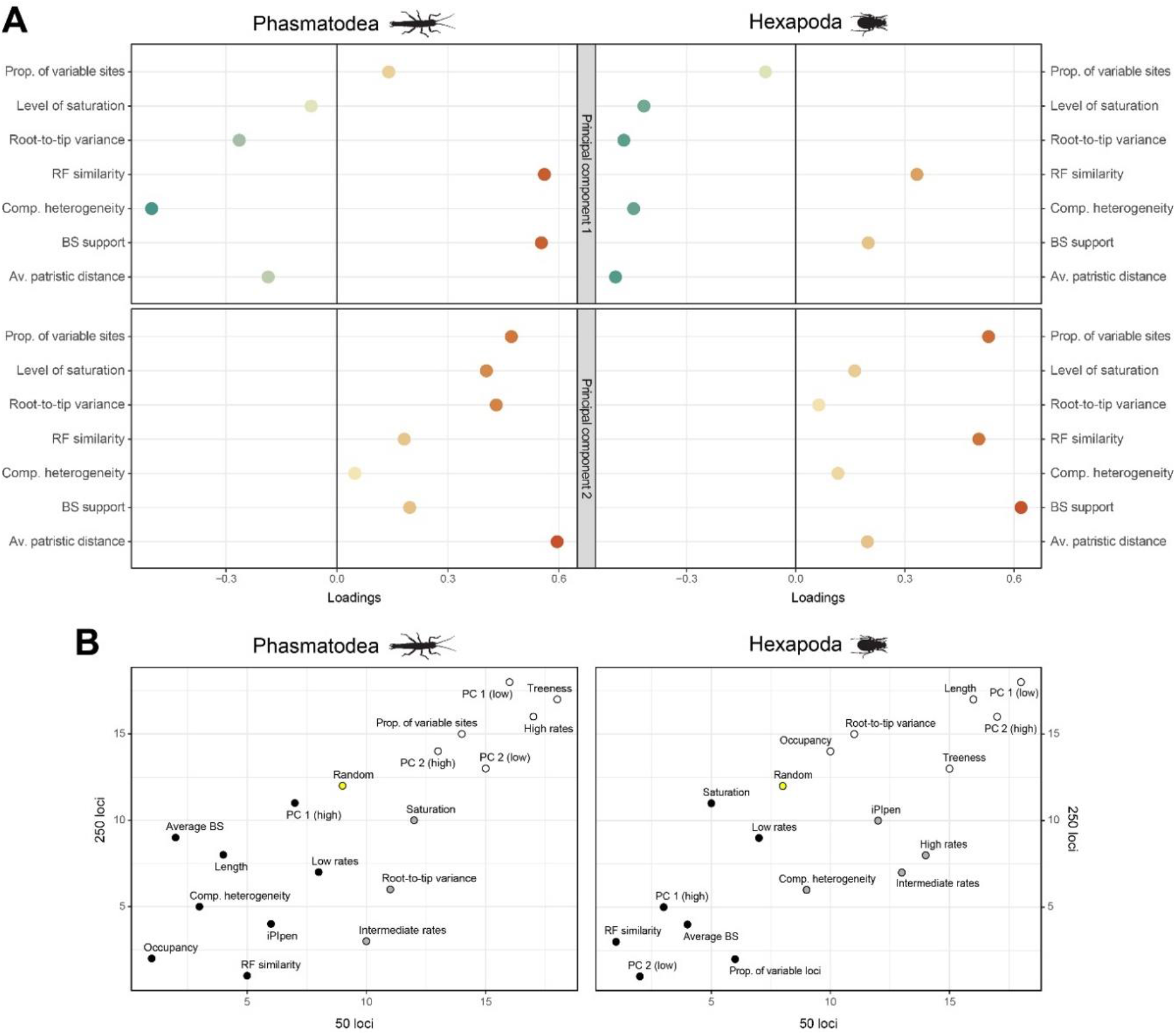
Results for the deviating datasets Phasmatodea (left) and Hexapoda (right). **A.** Loadings confirm the results of clustering analyses, showing that PC axes for these two datasets are reversed relative to those of the others, and that the signature of evolutionary rate is captured by principal component 2 (which shows all variables increasing/decreasing concomitantly), while that of usefulness is captured by principal component 1 (showing opposite trends for signal and bias). **B.** Ranking of subsampling strategies. Points in black rank higher than randomly sampled loci (yellow) for both 50 and 250 subsampling sizes, strategies shown in grey outperform random data only for 250 loci, and those in white rank lower for both. High usefulness (PC 1 for these datasets) ranks higher than random for both datasets and subsampling sizes. Including these results do not modify which strategies perform better than randomly-chosen loci at either subsampling size.

**Figure S7:**
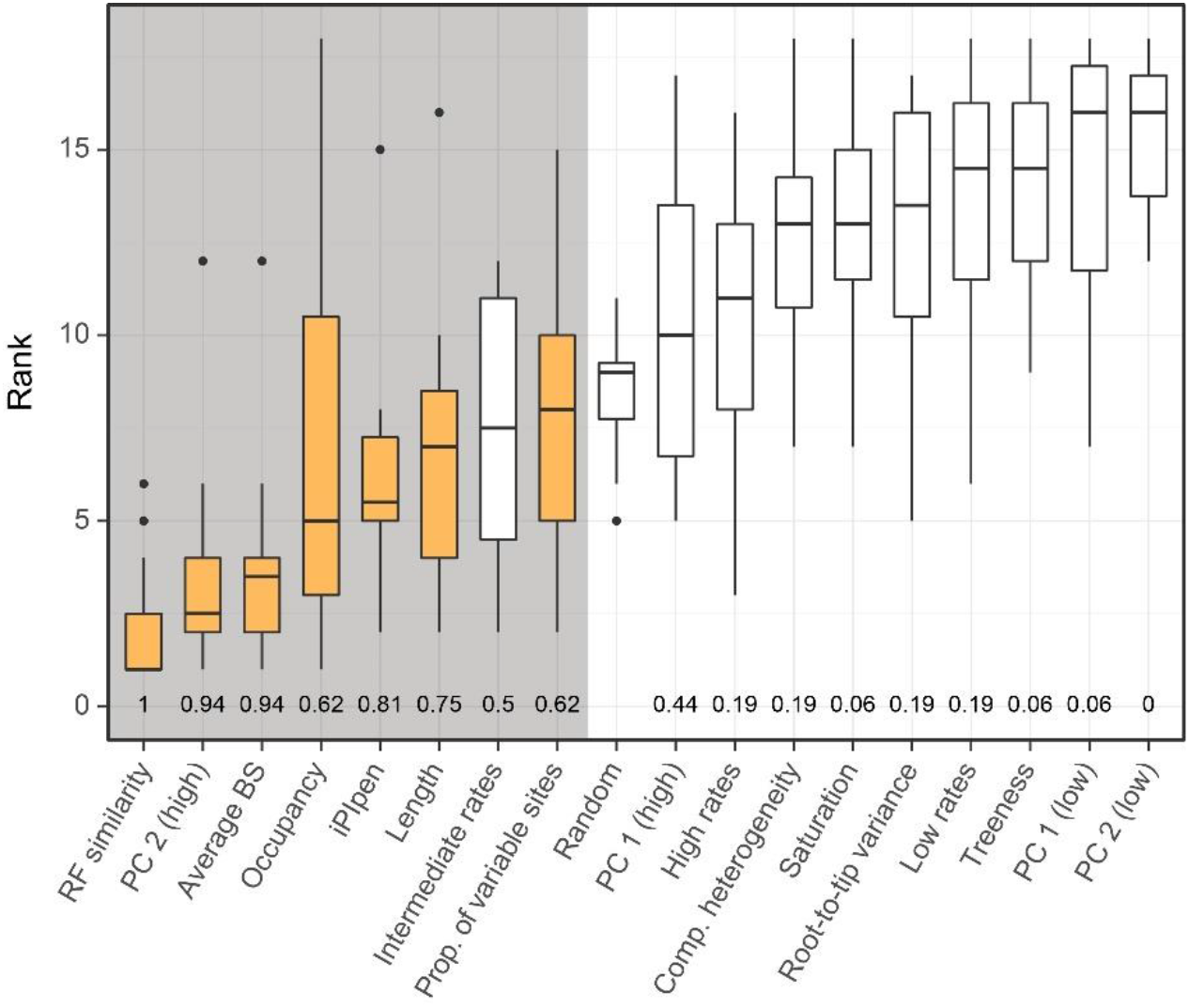
Performance of alternative strategies when subsampling to 50 loci. Overall, there is less variability in the ranks attained by strategies across datasets. Two criteria for selecting adequate strategies are highlighted: those whose median ranks are lower than randomly chosen loci (grey background), and those that outperform these in more than half of the datasets (yellow bars). The proportion of times a given strategy ranks lower than random loci is shown at the bottom. Results are very similar to those found for 250 loci (Fig. 3A), although both iPIpen and proportion of variable sites are also found here to outperform better than random loci selection across most datasets.

**Figure S8:**
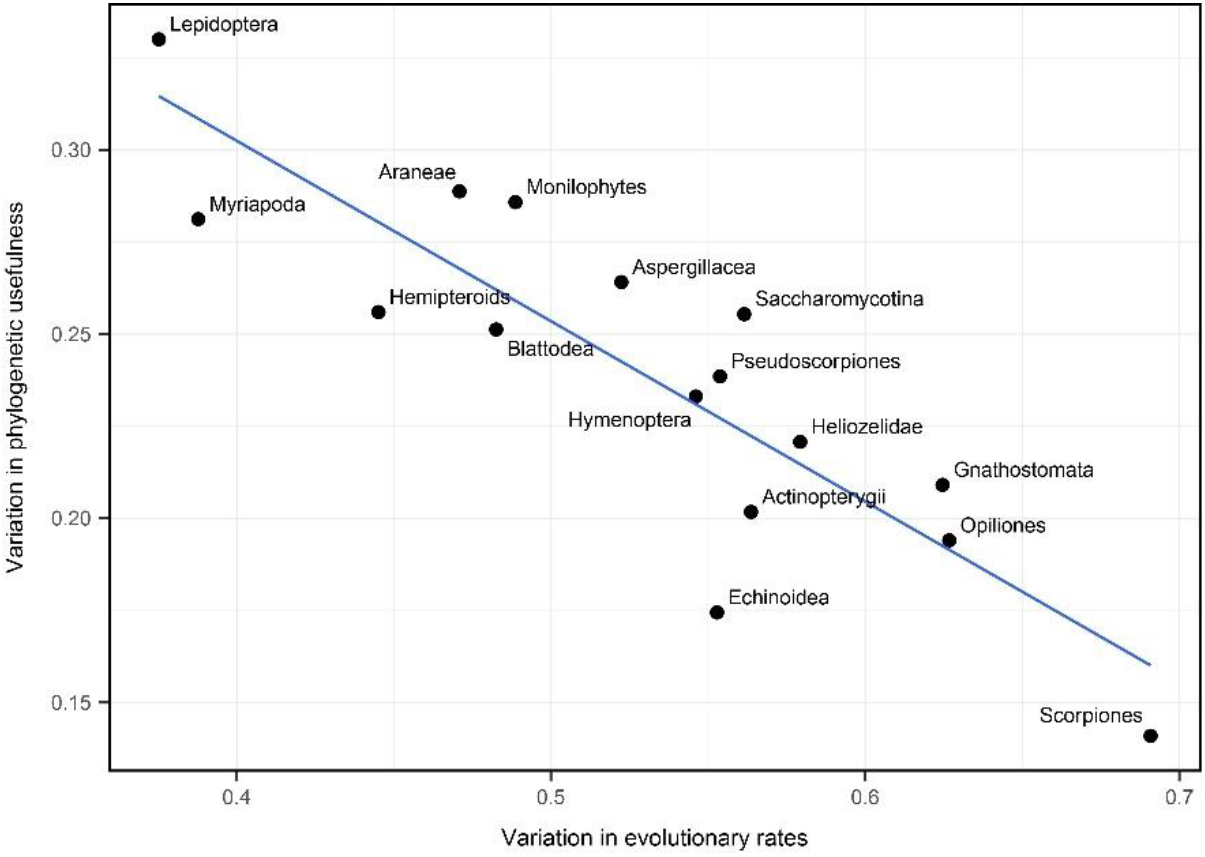
Phylogenomic datasets with large variations in evolutionary rates show less variation in phylogenetic usefulness. Axes correspond to the proportion of total variance explained by the first and second PC axes (interpreted for these datasets as capturing differences in rates and usefulness across loci, respectively). Correlation (ρ) between the two is −0.87, *P* = 1.2 × 10^−5^.

**Figure S9:**
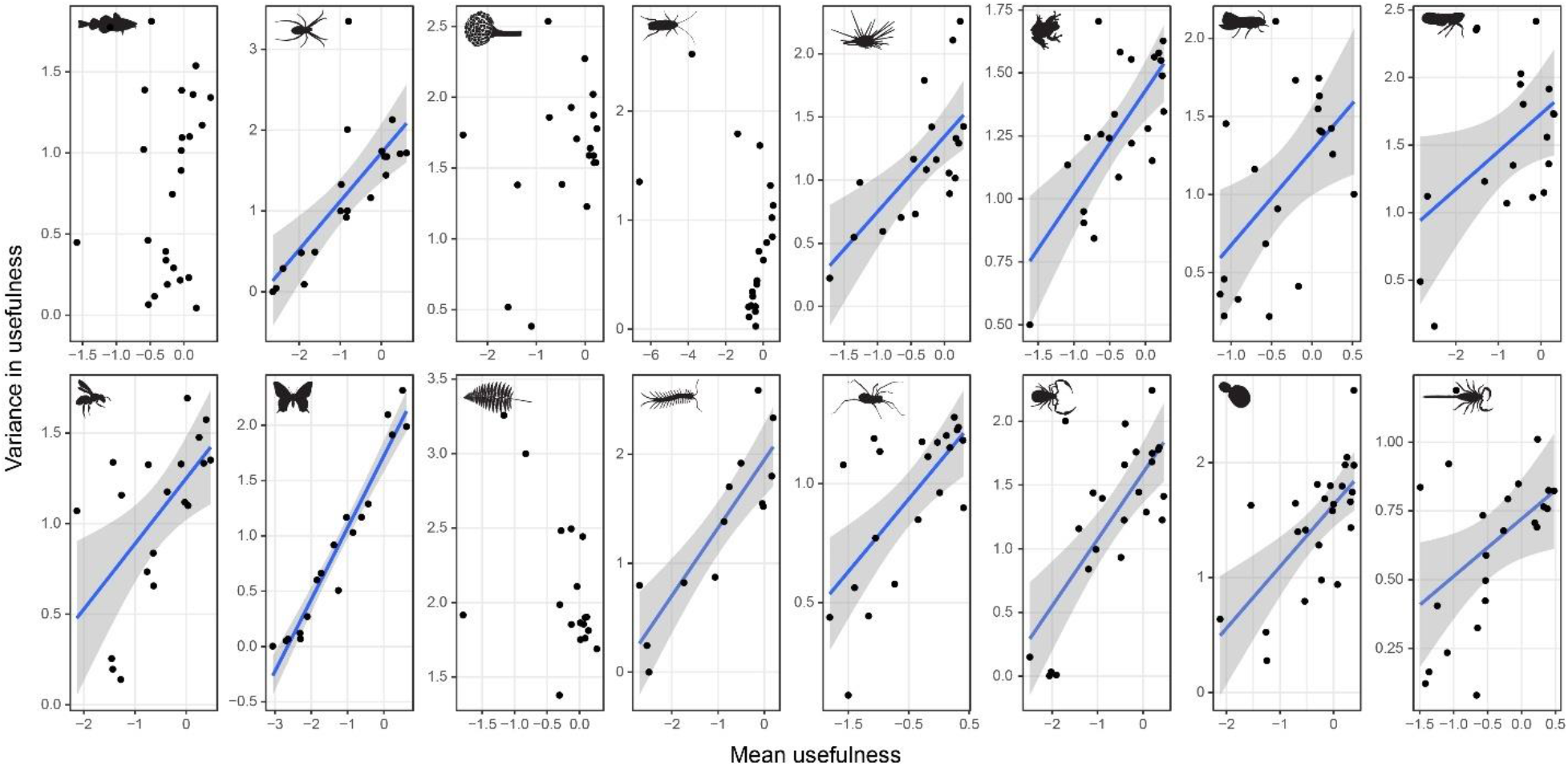
Genes evolving at optimal rates exhibit large differences in phylogenetic usefulness, such that many of them are still expected to perform poorly. Genes were classified into 25 categories obtained by subdividing tree-based evolutionary rates (log-transformed) into bins of equal width. The mean and variance of usefulness values (i.e., PC 2 scores) for loci within each bin was calculated, and fit to a linear regression (lines are shown only for clades with significant results).

